# Quantitative live imaging reveals region-dependent medulloblastoma dissemination in the zebrafish brain

**DOI:** 10.64898/2026.11.04.764295

**Authors:** Veronica Akle, Amin Allalou, Beyza Kilic, Sara Snyder, Joana Martins, Marc Thomas Schönholzer, Bernard Ciraulo, Martin Baumgartner, Stephan C.F. Neuhauss

## Abstract

Medulloblastoma is the most common malignant pediatric brain tumor, and relapse is frequently driven by dissemination within the central nervous system. Yet the early dynamics of tumor spread remain difficult to resolve *in vivo*. Here, we orthotopically transplanted human medulloblastoma cells into larval zebrafish and followed individual xenografts by longitudinal high-resolution confocal imaging. ONS-76 showed particularly robust dissemination, with hindbrain tumors spreading into the developing cerebellum in 81.5% of larvae, more than fourfold the frequency observed for forebrain or midbrain tumors. Disseminating cells extended along brain-surface and leptomeningeal-like regions and frequently associated with cerebral vasculature. Quantitative analysis of tumor area, perimeter, compactness, convex hull area, and fractal dimension captured changes in growth and spatial complexity over time. Dasatinib reduced tumor expansion and dissemination-related features in a concentration-dependent manner, while higher doses revealed host toxicity. These findings establish a powerful *in vivo* model for resolving medulloblastoma dissemination and therapeutic response.

## Introduction

Medulloblastoma is the most common malignant pediatric brain tumor and a leading cause of cancer-related mortality in children, owing to its rapid growth, its capacity to disseminate, and the morbidity associated with treating the developing nervous system [1,2]. Under the current World Health Organization classification, medulloblastomas comprise four molecularly distinct subgroups, WNT, SHH, Group 3 and Group 4, which differ in cell of origin, genetic alterations, metastatic propensity, and response to therapy [3].The behaviors that determine outcome, however, are functional rather than molecular: whether cells adhere, migrate, remodel their surroundings, and survive within brain tissue. These behaviors vary not only between subgroups but between individual tumors, and measuring them requires experimental systems in which the growth, dissemination and therapeutic response of a given sample can be read out directly, whether that sample is an established cell line, a primary culture, or patient-derived tumor cells. How closely such behaviors track subgroup identity, and how far they instead depend on the environment the cells encounter, remains poorly defined.

Dissemination and metastatic disease at diagnosis are associated with poorer survival and more complex treatment in medulloblastoma [4]. This increased risk is largely driven by the ability of tumor cells to spread within the central nervous system, where parenchymal invasion and leptomeningeal dissemination contribute to treatment resistance and subsequent relapse [5,6]. Accordingly, the presence and extent of dissemination are clinically important because they influence risk stratification and treatment intensity in patients with medulloblastoma.

The route and risk of spread also appear to depend on where the tumor arises: Group 3 and Group 4 tumors are located more frequently along the cerebellar midline, close to the ventricular system and cerebrospinal fluid pathways, whereas SHH tumors more commonly arise in the cerebellar hemispheres [7,8]. Whether disseminative behavior is dictated primarily by tumor-cell-intrinsic properties or by the anatomical compartment the cells occupy is difficult to establish, because dissemination in patients is inferred from imaging at diagnosis or from fixed material rather than observed as it unfolds.

Resolving this requires a setting in which tumor cells can be placed at a defined position within an intact brain and then followed over time. The brain microenvironment is not a passive substrate: extracellular matrix composition, resident neural and immune cells, the structure of the vasculature, and the architecture of the central nervous system, particularly the posterior fossa, together regulate tumor cell proliferation, migration, invasion, and response to treatment [9,10]. Orthotopic transplantation preserves many of these interactions, and orthotopic brain xenografts have established that human tumor cells engraft, invade and respond to treatment within a physiologically relevant neural microenvironment, in both glioblastoma and medulloblastoma [11–14].

Larval zebrafish are particularly well suited to this. During early development they lack a mature adaptive immune system, allowing engraftment of human tumor cells, including primary cultures and patient-derived material without immunosuppression [15–17]. Their optical transparency permits repeated, non-invasive imaging of the same animal, so that tumor growth, cell morphology and dissemination can be followed as a trajectory[18,19]. Their small size and compatibility with immersion-based drug exposure further allow therapeutic response and host tolerability to be examined in the same animals over short experimental periods. Building on these properties, zebrafish xenograft platforms combining standardized positioning with automated, high-content imaging have demonstrated that tumor progression can be followed *in vivo* at the scale required for compound screening [20,21]. In these approaches, tumor burden is typically quantified as fluorescence intensity or tumor area, measures that report reliably on how much tumor is present. The same images, however, also carry information about where the cells have gone: the extent of the territory a tumor occupies and the complexity of the boundary it forms. Extracting such descriptors alongside area descriptors would make dissemination itself a quantitative endpoint.

Here we used orthotopic larval zebrafish xenografts to ask how medulloblastoma cells spread within an intact vertebrate brain. We compared engraftment and dissemination across medulloblastoma cell lines, tested whether dissemination depends on the anatomical site of transplantation, and visualized the interactions of disseminating cells with the cerebral vasculature and with resident innate immune cells. Longitudinal imaging of individual xenografts was combined with measurements of tumor area, perimeter, compactness, convex hull area and fractal dimension, capturing changes in tumor burden, spatial extent and morphological complexity over time. We then applied this approach to dasatinib, a clinically approved inhibitor of BCR-ABL and SRC-family kinases, assessing its effects on tumor expansion, dissemination, proliferation and apoptosis alongside toxicity in the host. Because these readouts depend only on labelled tumor cells rather than on properties of a particular cell line, the approach could in principle also be applicable to primary cultures and patient-derived samples. It allows tumor morphology and progression to be investigated within an intact vertebrate central nervous system while retaining the scalability and imaging accessibility required for medium-throughput preclinical studies, from the assessment of therapeutic response in patient-derived material to the evaluation of novel compounds.

## RESULTS

To investigate medulloblastoma progression in vivo, we established an orthotopic larval zebrafish xenograft workflow combining brain transplantation, longitudinal imaging, and quantitative analysis of tumor morphology (Fig. 1a). Fluorescently labeled tumor cells were transplanted into the developing brain, and individual xenografts were repeatedly imaged to follow tumor progression over time. Segmented tumors were quantified using complementary morphometric parameters capturing changes in tumor size, spatial extent, and morphological complexity. This longitudinal approach was subsequently used to characterize tumor progression and treatment responses in vivo. The complete workflow is shown in Supplementary Fig. S1 and detailed in the Methods.

**Figure 1.**
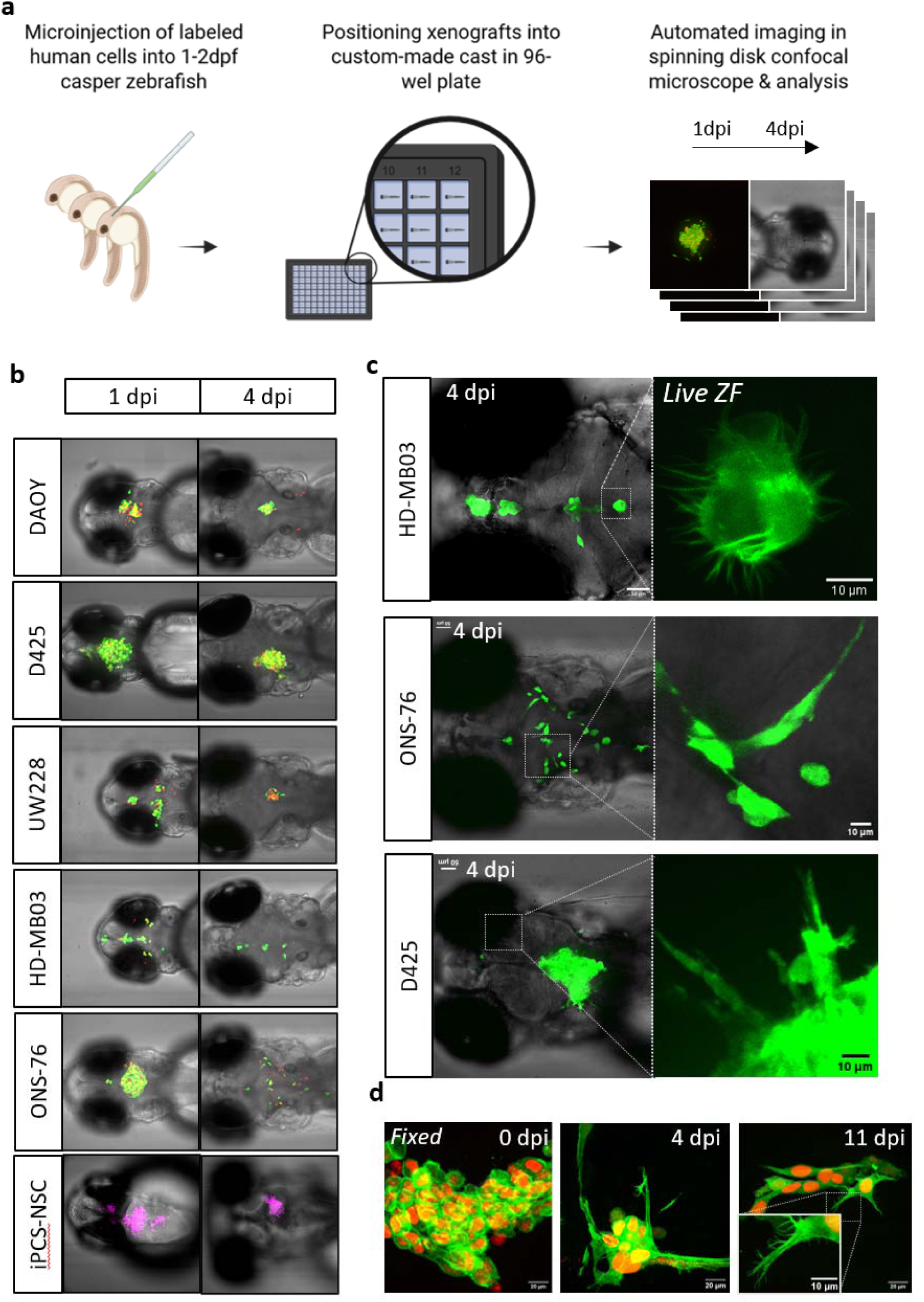
Comparison of xenotransplantation of medulloblastoma cells in brain of larva zebrafish. **a**. General workflow for longitudinal imaging of fluorescent tumor progression in brain of zebrafish. **b**. Tumor progression in the brain of zebrafish at 1 and 4 days post injection (dpi) of human medulloblastoma cell lines DAOY, D425, UW228, HD-MB03, ONS-76 and non-cancer-iPCS. **c**. Images of xenotransplants with different cells lines in live zebrafish at 4dpi. **d**. Confocal microscope 40X images of fixed xenotransplants with ONS-76 cells at 0, 4 and 11dpi. dpi= days post injection.

### Engraftment dynamics of medulloblastoma cell lines

To evaluate the suitability of the zebrafish larvae system to discriminate and study different cancer cells in the brain and other regions of zebrafish, we first compared the grafting capability of five commonly used medulloblastoma cell lines and one non-cancerous iPSC neural stem cell line. We used LA-eGFP mCherryNuc6 expressing ONS-76, HD-MB03, D425, DAOY and UW228 cell lines and fluorescently labeled iPSC neural stem cells. Approximately 300 cells were injected into 24 h post fertilization (hpf) *casper* larvae fish and maintained at 34 °C. Zebrafish larvae with xenografts were imaged using a spinning disk microscope at 1 and 4 days post injection (dpi) (Fig 1).

Following transplantation, medulloblastoma cells formed localized tumor masses within the developing brain. Live imaging revealed distinct patterns of tumor morphology and dissemination among the different cell lines. HD-MB03 cells frequently formed thin, filamentous protrusions, whereas ONS-76 and D425 cells displayed broader cellular extensions and more pronounced local spreading from the initial graft site (Fig. 1b). Approximately 90% of xenotransplanted larvae survived the first 24 h after injection (hpi), and over 80% remained viable through 4 dpi. A subset of ONS-76 xenografted larvae survived up to 11 dpi, with tumor cells still detectable in the brain; however, 4 dpi was selected as the experimental endpoint because dissemination was clearly established while larval survival remained high. Although the transplantation procedure produced xenografts of relatively similar initial size and anatomical location, tumor morphology and spatial distribution became increasingly variable between individual larvae over time (Fig. S2).

To compare the behavior of medulloblastoma cells to non-cancerous cells, iPCS-derived neural stem cells labeled with the lipophilic dye CellTrace® Deep Red (Thermo Fisher Scientific) were injected into the developing brain of larval zebrafish. These cells remained detectable for more than 24 h but displayed a morphology distinct from that of the medulloblastoma xenografts. They largely remained as compact clusters and extended long, thin cellular processes rather than showing the broader local spreading observed in the tumor cells (Fig. 1b; Supplementary Fig. S3).

For comparison, medulloblastoma cells were also transplanted into the yolk sac and perivitelline space. Xenografts formed at both sites, but these locations were less suitable for longitudinal analysis. Yolk-sac xenografts showed a rapid reduction in detectable tumor signal and limited optical accessibility, whereas perivitelline-space xenografts displayed variable morphology that depended strongly on the initial distribution of transplanted cells (Fig. S4). These observations supported the use of orthotopic brain xenografts for subsequent analyses of medulloblastoma progression and dissemination.

### ONS-76 medulloblastoma cells show region-dependent dissemination in the zebrafish brain

ONS-76 xenografts displayed different dissemination behavior depending on their location within the brain. Following transplantation into the upper rhombic lip or rhombencephalic ventricle, tumor masses were initially localized within the ventricular spaces of the forebrain, midbrain and hindbrain (Figs. 1–2). At 1 dpi, ONS-76 cells generally formed rounded tumor masses within the ventricles, although some cells had already begun extending into the surrounding brain parenchyma (Fig. 2a).

**Figure 2.**
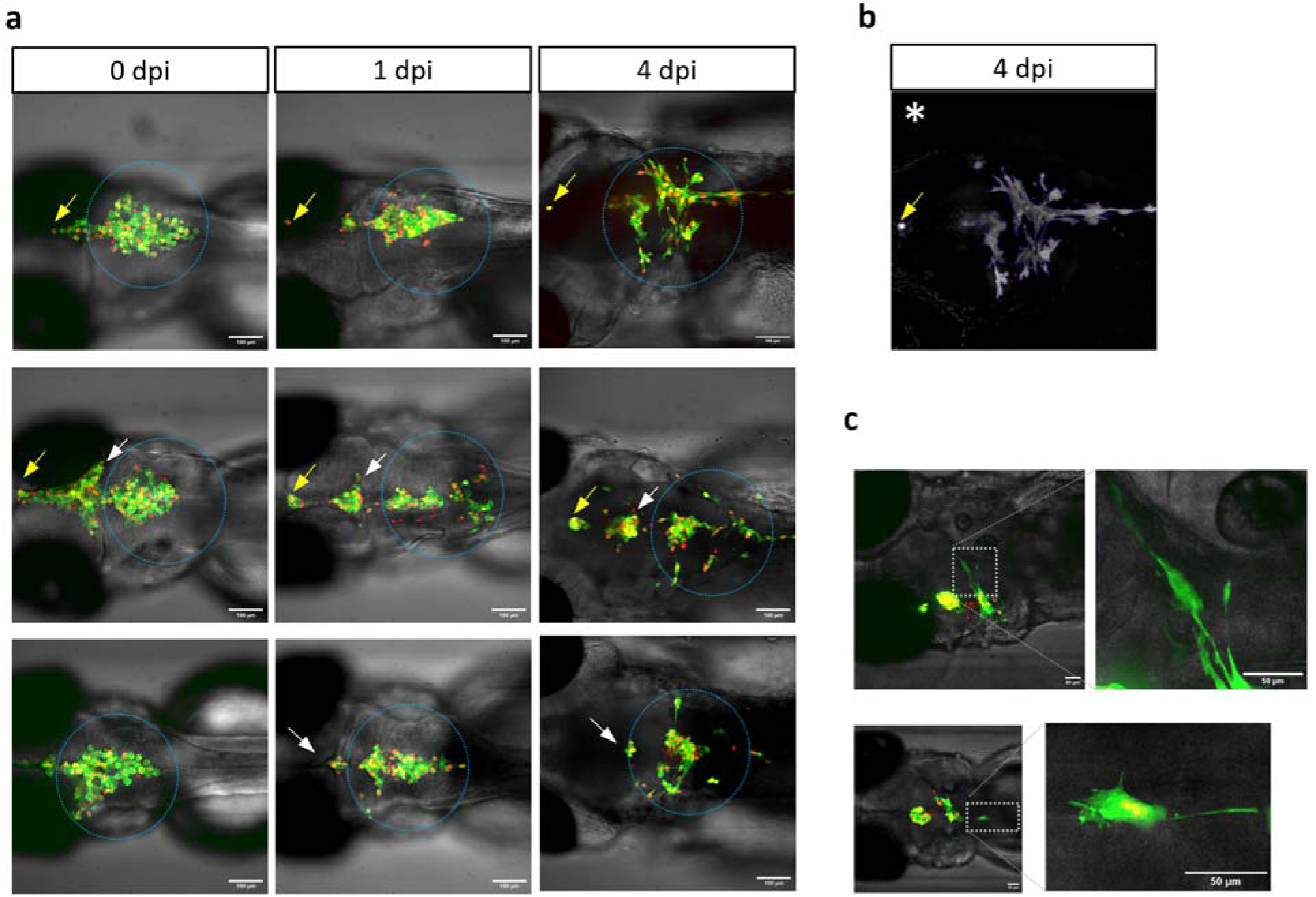
Medulloblastoma ONS-76 cells dissemination patterns. **a.** Tumor progression in three live zebrafish larvae at 0,1,4 dpi, showing that xenografted cells in the forebrain (yellow arrows remain small and compact or disappear, while cells in the midbrain (white arrows) disseminate to a lesser degree. On the contrary, cells in the hindbrain (blue circle) spread greater distances from the hindbrain after 4 dpi. **b**. Extracted maximal projection intensity image of the tumor example in a. **c.** Dissemination of tumor cells around the hindbrain along the leptomeninges at 4dpi.

This regional difference became particularly evident during subsequent tumor progression. ONS-76 cells located in the forebrain and midbrain ventricles invaded the adjacent tissue in only 20% of larvae (average of three independent experiments; n = 169 larvae). In the remaining cases, tumors persisted as relatively compact, small, rounded tumors with limited dissemination (white and yellow arrows in Fig. 2). In contrast, ONS-76 cells that localized to the hindbrain exhibited extensive spreading, with tumor cells disseminating into the developing cerebellum in 81.5% of larvae (average of three independent experiments; n = 189 larvae). These cells extended long cytoplasmic protrusions into the surrounding parenchyma and showed substantially more invasive morphology (Fig. 2 blue dotted area) than tumors located in anterior brain regions. In many hindbrain xenografts, individual tumor cells detached from the primary mass and migrated laterally along the brain surface and leptomeningeal-like regions, as well as caudally along the spinal cord, reaching regions distant from the injection site (Fig. 2c).

Together, these observations reveal a strong association between anatomical location and ONS-76 dissemination, with hindbrain/cerebellar xenografts showing a more than fourfold higher frequency of invasion than xenografts located in the forebrain or midbrain. This pronounced regional difference suggests that local features of the hindbrain/cerebellar environment may contribute to the invasive behavior of ONS-76 cells.

### Relationship of the tumor with microenvironment

To investigate the relationship between medulloblastoma cells and the vascular microenvironment, LA-iRFP-labeled ONS-76 cells were orthotopically transplanted into the brains of *Tg(kdrl:mCherry)* zebrafish larvae, in which endothelial cells are fluorescently labeled with GFP (Fig. 3). At 1 dpi, the tumor masses formed in close proximity to the cerebral vasculature without inducing obvious alterations to the vascular architecture. Confocal imaging revealed that the developing blood vessels remained largely intact as tumor cells expanded around them, suggesting that early tumor growth occurs within and adapts to the developing vascular network.

**Figure 3.**
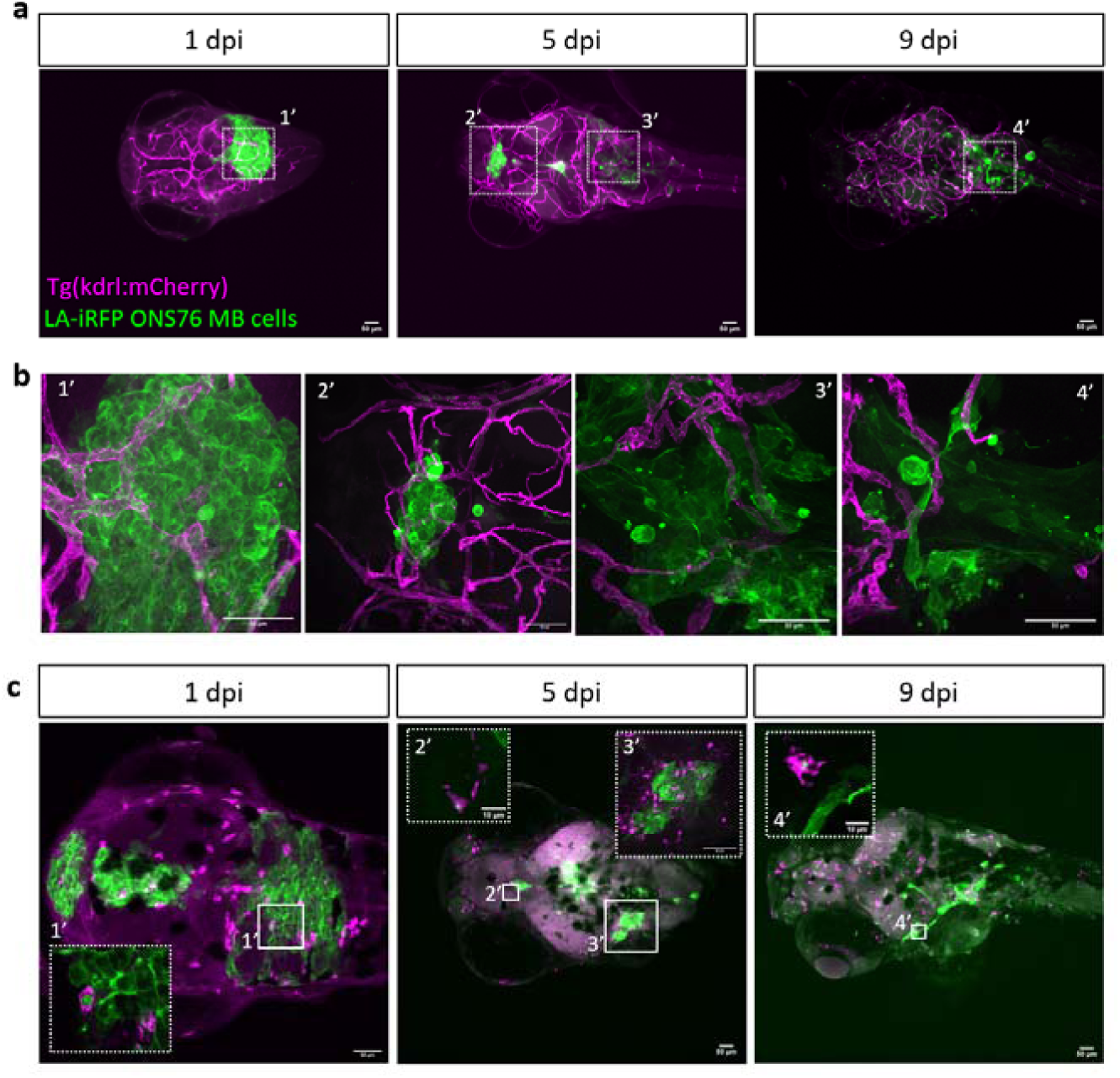
Medulloblastoma ONS-76 cells interact with the vascular and innate immune microenvironment. **a.** Representative *Tg(kdrl:mCherry)* zebrafish larvae at 1, 5, and 9 dpi showing LA-iRFP-labeled ONS-76 xenografts and the cerebral vasculature (magenta). **b**. High magnification confocal images (40x) of selected regions in a. **c.** Representative *Tg(mpeg1:mCherry)* larvae at 1, 5, and 9 dpi, showing the relationship between disseminating ONS-76 cells (green) and microglia. Inserts show engulfment of tumor cells/fragments by macrophages (1′-4′).

At 5 and 9 dpi, disseminating ONS-76 cells showed a pronounced spatial association with cerebral blood vessels, particularly in the hindbrain. Tumor cells were frequently observed along the trajectory of individual vessels and at sites distant from the primary tumor mass, while remaining closely associated with the vascular network. Three-dimensional reconstruction further showed tumor cells extending around and along blood vessels during dissemination (Supplementary Video 1). Together, these observations reveal a persistent association between disseminating ONS-76 cells and the cerebral vasculature.

We next examined interactions with innate immune cells by transplanting LA-iRFP-labeled ONS-76 cells into *Tg(mpeg1:mCherry)* zebrafish larvae. As early as 1 dpi, numerous macrophages/microglia were present throughout the brain and in direct contact with the xenografts (Fig 3c). These cells were frequently observed at the tumor border and interspersed among tumor cells, although they remained broadly distributed throughout the brain and did not show an obvious preferential accumulation around the xenograft. At later time points, macrophages/microglia remained closely associated with the xenograft and were observed both within the tumor mass and in the surrounding brain parenchyma (Fig. 3). Tumor-derived fluorescent material was occasionally observed within GFP-positive cells, and individual tumor cells or fragments were occasionally seen closely associated with these cells, consistent with sustained interactions and uptake of tumor-derived material (Fig. 3c).

Quantitative analysis revealed that the total number of GFP-positive cells within the brain remained relatively stable between 1 and 5 dpi, with averages of 97.9 (n=5) and 102.1 (n=8) cells per brain, respectively, before declining to an average of 47.5 cells (n=7) by 9 dpi. Despite this overall decline, the proportion of macrophages/microglia associated with the tumor remained remarkably constant throughout the experiment, representing 29.98% of the total microglial population at 1 dpi, 23.97 % at 5 dpi, and 24.08 % at 9 dpi. Thus, the relative association of the tumor with the macrophage/microglia-lineage cells was maintained over time, indicating a persistent interaction, rather than progressive recruitment or accumulation around the xenograft.

### *In vivo* toxicity evaluation of dasatinib

To test whether the longitudinal xenograft assay could detect pharmacological modulation of tumor progression, we evaluated dasatinib, a clinically approved tyrosine kinase inhibitor targeting BCR/ABL, SRC-family kinases, c-Kit, Eph receptors, and other tyrosine kinases. Before evaluating tumor responses in the orthotopic model, we established a tolerated concentration range in zebrafish larvae.

Concentration-response experiments were performed to assess dasatinib-associated toxicity. Larvae were exposed to 5, 10, 15, 20 and 30 µM dasatinib and monitored for survival and developmental abnormalities. In parallel, automated VAST imaging was used to assess treatment-associated morphological effects, including alterations in body length, edema formation, spinal curvature, and overall developmental progression. Based on survival, the LC50 for dasatinib was 30 µM. However, morphometric analysis evidenced major effects of dasatinib already above 10 µM (Fig. 4a). Based on these observations, concentrations up to 15 µM were selected for subsequent xenotransplantation experiments.

**Figure 4.**
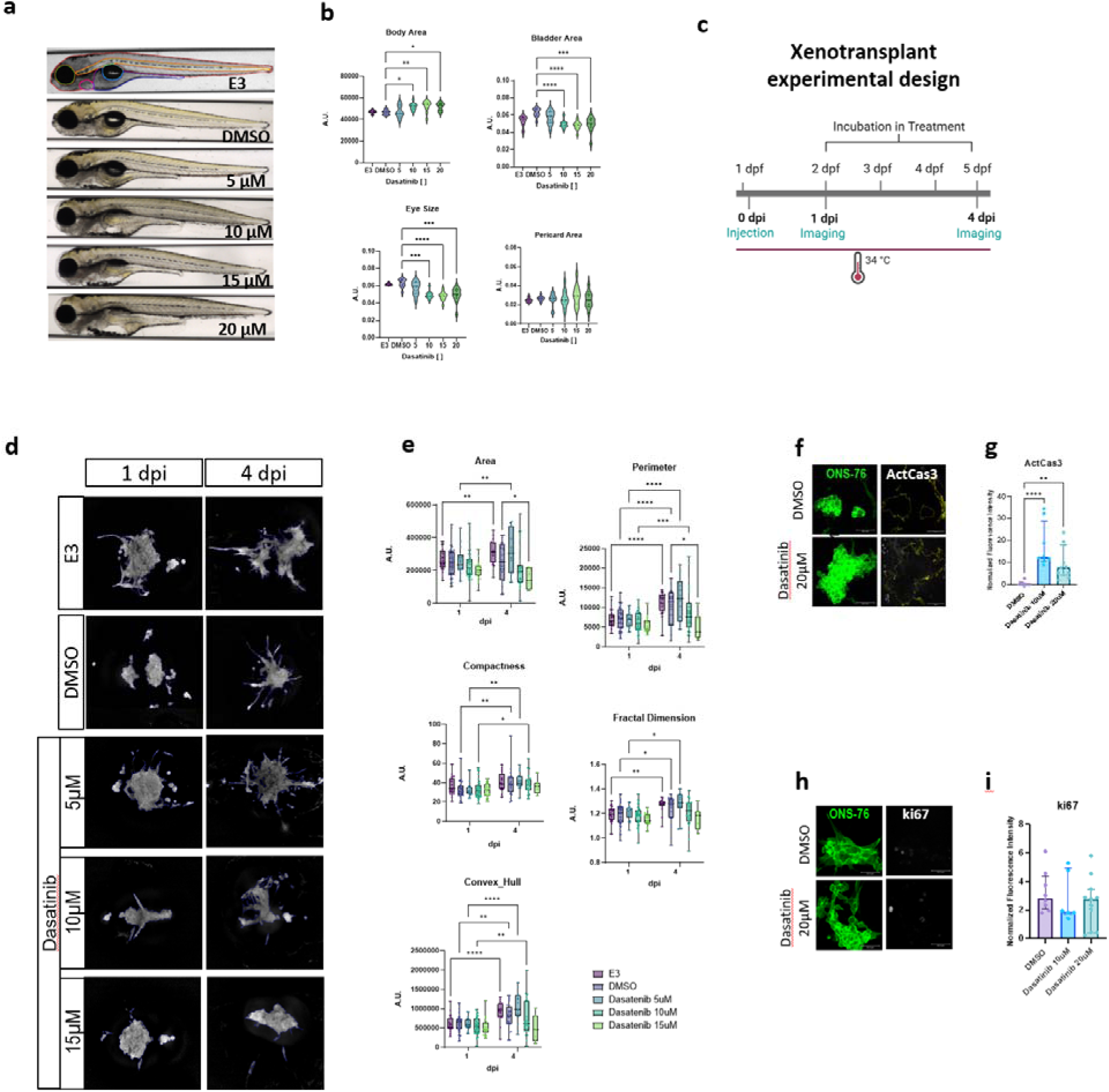
Effect of dasatinib in wild-type zebrafish embryos. **a**. Representative images of 5 dpf wild-type larvae treated with dasatinib at different concentrations for 3 days. **b**. Quantification of bod structures after dasatinib treatment. Kruskal-Wallis Test wit Dunn’s multiple comparisons test with DMSO as control, n=12 per group. **c**. Experimental design for the xenotransplant experiment. d. Representative maximum-intensity projections of same larvae with segmented ONS-76 xenografts imaged before (1 dpi; 48 hpf) and after 4 days of continuous treatment. **e**. Quantification of tumor morphological features: perimeter, area, compactness, fractal dimension, and convex hull area. Data analyzed using a mixed-effects model (REML) with repeated measures, followed by Tukey’s multiple-comparisons test. Individual larvae are shown as dots; boxes represent the median and interquartile range, and whiskers indicate the minimum and maximum values. P values are indicated in the graphs. **f**. Immunofluorescence for ActCasp3 in LA:GFP Nuc6:mCherry ONS-76 medulloblastoma cells at 4dpi. **g**. Normalized fluorescent intensity quantification of f. **h**. Immunofluorescence for Ki67 in LA:GFP Nuc6:mCherry ONS-76 medulloblastoma cells at 4dpi. **i**. Normalized fluorescent intensity quantification of h. The results are shown as median with interquartile range. Kruskal–Wallis test followed by Dunn’s multiple-comparisons test p < 0.05. Rostral to the left.

### Dasatinib restricts ONS-76 xenograft dissemination *in vivo*

Approximately 300–400 LA:eGFP-Nuc:mCherry ONS-76 cells were transplanted into the hindbrain of 24 hpf casper zebrafish larvae. Larvae with brain-localized xenografts were imaged at 1 dpi before treatment and re-imaged at 4 dpi following continuous exposure to 5, 10, or 15 µM dasatinib, DMSO vehicle, or E3 medium (Fig. 4c,d). This longitudinal design allowed changes in individual tumors to be followed over time using area, perimeter, compactness, convex hull area, and fractal dimension as complementary measures of tumor growth, spatial expansion, and morphological complexity.

Longitudinal analysis revealed a concentration-dependent inhibitory effect of dasatinib on tumor progression and dissemination (Fig. 4e). In E3 and DMSO-treated xenografts, tumor area, perimeter, and convex hull area increased significantly between 1 and 4 dpi (Supplementary Table 1), consistent with continued tumor expansion and spreading within the brain. In contrast, this increase was progressively attenuated in dasatinib-treated tumors, particularly at 10 and 15 µM. Mixed-effects analysis identified significant main effects of treatment on area (p = 0.0011), perimeter (p = 0.0017), convex hull area (p = 0.0067), and fractal dimension (p = 0.0138), together with significant treatment × time interactions for area (p = 0.0031), perimeter (p = 0.0155), and convex hull area (p = 0.0041), indicating that dasatinib altered the progression of these features over time.

Perimeter and convex hull area, which reflect extension and spatial distribution of the tumor, respectively, increased markedly in control xenografts but were reduced in a dose-dependent manner with higher dasatinib concentrations Similarly, tumor area expanded over time in control animals (E3, p=0.0022), whereas this increase was reduced with increasing dasatinib concentration. Across the four responding parameters, values at 4 dpi decreased progressively with increasing dasatinib concentration; in post-hoc comparisons at 4 dpi, the difference from DMSO reached significance at 15 µM for area (p = 0.0481) and perimeter (p = 0.0202). Fractal dimension increased over time in control and 5 µM xenografts (p = 0.0226) and was lower overall in dasatinib-treated animals, consistent with reduced morphological complexity, although the treatment × time interaction was not significant (p = 0.4088). In contrast, compactness increased significantly over time (p < 0.0001) but showed neither a treatment effect (p = 0.6986) nor a treatment × time interaction (p = 0.7365), indicating that the predominant effect of dasatinib was on tumor expansion and spatial dissemination rather than on this aspect of tumor shape.

To determine whether these morphological effects were accompanied by changes in tumor-cell proliferation or apoptosis, xenografts were analyzed at 4 dpi by immunofluorescence for Ki67 and activated caspase-3 (Fig. 4f–i). Ki67 fluorescence did not differ significantly between control and dasatinib-treated xenografts. In contrast, activated caspase-3 fluorescence within the tumor region was significantly increased following dasatinib treatment, indicating an increase in apoptotic signaling associated with treatment.

Together, these findings show that dasatinib restricts the longitudinal expansion and dissemination of ONS-76 xenografts *in vivo* and is associated with increased apoptotic signaling without a detectable reduction in Ki67 labeling.

## Discussion

Orthotopic transplantation of human medulloblastoma cells into the larval zebrafish brain revealed distinct patterns of tumor growth and dissemination within the first days after injection. Importantly, the optical transparency of zebrafish larvae allowed tumor cells to be visualized at high resolution within a living vertebrate brain. Thus, the relevant endpoints were not limited to tumor shrinkage but included cell dispersion, migration, changes in tumor shape, association with host structures, and nuclear morphology. Longitudinal imaging of the same animals further allowed these behaviors to be followed over time, providing information on spatial organization and tumor complexity that is difficult to obtain in less optically accessible *in vivo* models.

All medulloblastoma cell lines tested formed grafts and survived for at least 4 days, and some up to 11 days, although their subsequent behavior differed. These differences did not follow molecular subgroup classification. ONS-76, DAOY, and UW228 are classified as SHH models, whereas D425 and HD-MB03 are Group 3 models [22]. Despite both belonging to Group 3, D425 and HD-MB03 behaved differently *in vivo*: D425 cells were highly motile and disseminated extensively, whereas HD-MB03 formed more compact grafts. Previous studies have similarly shown differences in proliferation, migration, and invasion among medulloblastoma models, including in 3D brain-mimetic extracellular matrices. However, because the panel tested here was small and functional differences among the cell lines were not examined in depth, subsequent analyses focused primarily on ONS-76, which showed the most reproducible dissemination phenotype.

ONS-76 xenografts showed robust and reproducible spread from the injection site, with more than 80% of engrafted tumors disseminating widely into the hindbrain. This phenotype is particularly relevant because dissemination within the central nervous system is a major determinant of recurrence and poor outcome in medulloblastoma. The reproducibility and rapid onset of this behavior make ONS-76 a useful model for studying early dissemination and testing interventions that restrict tumor spread.

The disseminative behavior of ONS-76 may relate to the reported biological properties of the cell line. ONS-76 has been described as a primitive SHH medulloblastoma model with immature or stem-like features and high proliferative capacity [23], and can express neural progenitor or stemness-associated markers and show strong growth potential compared with other medulloblastoma lines [24,25]. Nevertheless, the mechanisms driving its dissemination remain unresolved and are likely to involve a combination of intrinsic migratory programs and interactions with surrounding brain structures.

A striking feature of ONS-76 xenografts was the route followed by disseminating cells. In addition to expanding into adjacent brain tissue, cells frequently extended away from the injection site along the surface of the brain and leptomeningeal-like regions. This is relevant because medulloblastoma dissemination in patients occurs largely through cerebrospinal fluid pathways and it is associated with leptomeningeal seeding, a major contributor to recurrence and poor outcome.

The location of transplantation likely contributes to this phenotype: injection into the rhombencephalic ventricle or adjacent hindbrain places tumor cells close to ventricular and brain-surface compartments that may provide permissive spaces for spread. Conceptually, this resembles intracisternal mouse models and other zebrafish models in which tumor cells introduced into ventricular or cerebrospinal fluid-associated spaces disseminate along brain-surface routes[26–28].

Primary tumor location may also influence dissemination: Group 3 and Group 4 tumors more frequently arise near the cerebellar midline and ventricular system, whereas SHH tumors more commonly arise in the cerebellar hemispheres [29,30]. Although ONS-76 is an SHH model, its reproducible spread in the zebrafish hindbrain indicates that the local transplantation environment can reveal disseminative potential when tumor cells are placed near permissive anatomical routes. Conversely, site selection should be considered a design variable rather than a technical detail.

The spatial pattern of spread suggests that ONS-76 cells may use more than one anatomical route. Co-imaging with the vascular reporter *Tg(kdrl:mCherry)* showed tumor cells frequently associated with vessels at the tumor periphery, with some cells appearing to wrap around vascular structures (sup. Video). These qualitative observations suggest that vascular surfaces may provide adhesion sites, provide growth cues or guide local extension. At the same time, cells may disperse through ventricular or brain-surface spaces. Distinguishing patterns and modes of active migration along host structures and permissive anatomical compartments will require time-lapse imaging at higher temporal resolution.

Cells located away from the main xenograft also frequently displayed oval or elongated nuclei. Nuclear deformation may reflect mechanical confinement during migration through narrow anatomical spaces [31,32]. In melanoma, mechanical confinement deforms nuclei and promotes a phenotypic switch from a proliferative toward a more invasive state [33]. Our data do not establish an equivalent mechanism in medulloblastoma, but the nuclear morphology of disseminated ONS-76 cells raises the possibility that migrating cells transiently adopt a less proliferative and more invasive state. Combining nuclear-shape analysis with proliferation markers could test this hypothesis.

Microglia reporter larvae allowed early interactions between tumor cells and the innate immune environment to be visualized. Microglia were present throughout the brain and at the tumor borders, but we did not observe increased accumulation within or immediately surrounding ONS-76 xenografts.

ONS-76 xenografts therefore did not induce strong microglial recruitment during the early period examined. This resembles zebrafish glioblastoma xenografts, where microglia interact with tumor cells without extensive engulfment [34]. Whether these interactions support tumor growth or primarily reflect debris clearance remains to be tested. Microglial numbers followed the expected developmental increase between 3 and 5 dpf [35], although functional activation without a change in cell number cannot be excluded.

Dasatinib treatment restrained the expansion of ONS-76 xenografts. Tumor area, perimeter, and convex hull area increased less under treatment, and fractal dimension was lower in treated animals, whereas compactness was not significantly altered. Together, these parameters indicate that dasatinib affected not only tumor size but also the extent and complexity of tumor spread. The concentration-dependent changes across several morphometric parameters further support a biological effect on tumor expansion.

Analysis of tumor-cell state indicated that the reduction in tumor burden was associated primarily with increased apoptosis rather than reduced proliferation. Cleaved caspase-3 staining increased after dasatinib treatment, whereas proliferation was not detectably altered. Dasatinib also targets additional receptor tyrosine kinases and has variable effects on proliferation while inducing apoptosis in a cell line-dependent manner and consistently disrupting adhesion, migration, and invasion [36]. Thus, the reduction in tumor area observed here appears to reflect increased cell death rather than cell-cycle arrest. The morphological response is consistent with previous *in vitro* studies showing that dasatinib interferes with pathways involved in medulloblastoma cell motility. Here, reduced spatial expansion and increased tumor-cell death occurred within the same xenograft, illustrating the value of measuring dissemination in addition to tumor burden.

Dasatinib did not cause mortality at the concentrations tested, but concentrations above 10 µM produced generalized edema, indicating developmental toxicity. The strongest anti-tumor effects therefore occurred near the threshold at which the host was also affected. This limits the interpretation of dasatinib as a selectively effective treatment under the present exposure conditions but also illustrates an important advantage of the model: anti-tumor efficacy and organismal tolerability can be assessed simultaneously.

Combination approaches are also worth pursuing. Dasatinib synergizes with cisplatin in medulloblastoma cells through combined SRC and STAT3 inhibition[37], while combined SRC and aurora kinase inhibition markedly enhances suppression of medulloblastoma survival and migration compared with either agent alone [38]. Testing such combinations in the zebrafish xenograft system could determine whether anti-tumor activity can be increased at lower, better-tolerated dasatinib concentrations.

### Limitations and outlook

Several limitations should be considered. Morphometric parameters were derived from two-dimensional projections and therefore approximate the three-dimensional organization of the graft; volumetric imaging would provide a more complete description but reduce throughput. Larvae were maintained at 34°C, which may influence both host and tumor-cell behavior.

In addition, the 4-day observation window restricts the analysis to early events after transplantation, so phenotypes requiring longer adaptation to the brain microenvironment, such as delayed dissemination, may be missed. The zebrafish brain also differs from the human cerebellum in cytoarchitecture, extracellular matrix composition, and meningeal organization; therefore, the spread described here should be regarded as anatomically analogous rather than equivalent to leptomeningeal dissemination in patients. Incompatibilities between human receptors and zebrafish ligands may further limit some microenvironmental interactions.

## Conclusion

Overall, this platform serves as a bridge between *in vitro* drug screening and more complex *in vivo* models. In cell culture, compound activity is usually interpreted through direct effects on tumor-cell viability, proliferation, or migration. In the larval xenograft system, these readouts can be assessed alongside tumor morphology, dissemination, apoptosis, and organismal toxicity. This is particularly relevant for early drug-development studies, including the evaluation of new compounds or combination therapies, where both efficacy and tolerability must be considered before advancing to more complex animal models.

## METHODS

### Cell culture

ONS-76 (SHH MB, TP53 WT, NRAS 128A>G mutation) cells were generously provided by Michael Taylor (SickKids, Canada) and HD-MB03 MB03 (Non-WNT/non-SHH MB, TP53 mutant, MYC amplification) cells were generously provided by Till Milde (DKFZ, Germany) and both cultured in Roswell Park Memorial Institute 1640 medium (RPMI, Gibco). D425 (Group 3 MB, TP53 mutant, MYC amplification) cells were generously provided by the Henry Friedman lab (Duke University, UK) and cultured in Improved Modified Eagle Medium (IMEM, Gibco). UW228 (SHH MB, TP53 mutant) cells were generously provided by John Silber (Seattle, USA) and cultured in Dulbecco’s Modified Eagle Medium (DMEM, Sigma). DAOY (SHH MB, TP53 WT) cells were purchased from the American Type Culture Collection (ATCC, Rockville, MD, USA) and cultured in Iscove’s Modified Dulbecco’s Medium (IMEM, Sigma). All labeled cell lines were produced by lentiviral transduction of wildtype ONS-76, D425, HD-MB03, UW228 and DAOY cells with pLenti-LA-EGFP, mCherryNuc, or LA-iRFP. iPSC-derived neural stem cells were generously provided by Ruxandra Bachmann. The cells were raised at 37°C in a humidified atmosphere containing 5% CO2 and then adapted to 34°C, 2% CO_2_ environment. LookOut® Mycoplasma PCR Detection kit (#MP0035, Sigma Aldrich) and MycoAlert (Lonza, Visp, Switzerland) were used to exclude mycoplasma contaminations of cell lines used. All cell lines were authenticated by Single Nucleotide Polymorphism (SNP)- Profiling.

### Animals and housing conditions

Adult zebrafish of the AB and *roy-/- nacre -/-* (casper) strains were housed in the Zebrafish Facility at Zurich University, under optimal conditions for the species. They were maintained in a multi-tank recirculating water system under a 14-hour light/10-hour dark cycle, with a temperature of 28 (±1 °C), a salinity range of 600–800 S/cm, and a pH between 7.0 and 7.5. Embryos were obtained by natural spawning and raised at 28.5 C in Embryo Water (E3) until needed. All animal experiments were conducted in accordance with applicable institutional and national regulations. All protocols and experimental procedures involving zebrafish were approved by the Cantonal Veterinary Office, Zürich (Kantonales Veterinäramt), under licenses. All animal experiments were conducted at the University of Zurich in accordance with Swiss animal welfare regulations and under authorization from the Veterinary Office of the Canton of Zurich (license no. TV4206 and ZH073/2024). The Cantonal Commission for Animal Experimentation serves as the independent ethical review body, while the University of Zurich oversees institutional compliance through its Animal Welfare Officers.

### Toxicity Assessment

At 2 days post fertilization (dpf), dechorionated wildtype AB embryos were exposed to dasatinib (Sigma-Aldrich, CDS023389) at 1–40 µM; n=25/group or controls (E3 or 0.15% DMSO) to determine LC50. Larvae were monitored every day and deaths were registered. At 5 dpf, larvae were imaged using the Vertebrate Automated Screening Technology (VAST) BioImager platform for morphometric analysis. . A ResNet50-based segmentation model, trained on 2,250 annotated images, was used to quantify body area, eye size, yolk sac area, and swim bladder inflation across all experimental images[39]. Lateral views were used for quantification. The concentrations that did not produce deaths were used for xenograft treatments.

### Xenotransplantation

On the day of injection, cells were collected using TrypLE®, centrifuged and filtered through a 40µm cell strainer to reduce cell clumping. About 2×10^6^ cells were resuspended in 40 µl of 1% polyvinylpyrrolidone (PVP) in cell culture media, which increased viscosity and prevented cell aggregation during microinjection. Tumor cells were kept on ice and microinjected within 1h after collection.

### Orthotopic xenotransplantation and larval selection

Casper embryos were collected and maintained at 31 °C. At 24 hours post fertilization (hpf), embryos were manually dechorionated, anesthetized, and placed in agarose-lined Petri dishes for microinjection. Approximately 300–400 tumor cells were injected into the ventricular space of the developing hindbrain. Injected larvae were maintained at 34 °C until the following day. At 48 hpf (1 day post injection, dpi), larvae were screened for fluorescent microtumors confined to the brain. Larvae with tumor cells in other regions of the head or body were excluded from the study. Tumor-bearing larvae were anesthetized with 0.2 mg/mL tricaine (MS-222) before positioning and imaging.

### Custom agarose molds and longitudinal positioning

To enable reproducible dorsal brain imaging of individual larvae in an inverted confocal microscope, custom 3D-printed inserts were designed to fit square 96-well ibidi plates. Each insert consisted of a linear array of eight molds corresponding to one plate column. Each mold had a three-level profile: a rectangular base fitting the well, a middle reservoir that allowed solution exchange, and a narrow ridge that generated a confined space in which the larva could be passively restrained while remaining submerged. All levels were tapered to facilitate demolding. Agarose (1.5% w/v) was prepared in E3 medium and used to cast the positioning cavities in the plate.

The molded cavities positioned larvae ventral side down (supine), bringing the dorsal brain close to the glass bottom of the plate and providing a consistent orientation for repeated imaging. At later stages, when larvae were thinner, 0.08% agarose containing 2 mg/mL tricaine methanesulfonate (MS-222) in E3 was used to stabilize the larvae and minimize tilt during imaging. The 3D mold master was fabricated in BioMed Clear Resin (Formlabs), selected for its smooth surface and ease of demolding. Design STEP files and dimensional specifications are provided in the Supplementary Information.

### Longitudinal image acquisition

Tumor-bearing larvae were positioned dorsal side down, one per well, in the custom agarose molds. Imaging was performed with an Olympus IXplore SpinSR10 spinning-disk confocal microscope equipped with a Yokogawa CSU-W1 50 µm pinhole disk, a long-working-distance UPLXAPO 20× air objective, and a Hamamatsu ORCA-Fusion sCMOS camera (2304 × 2304 pixels; 6.5 × 6.5 µm pixel size). A 4× bright-field overview image was acquired first. Larval heads were located using a trained detection algorithm implemented in cellSens Dimension software, and each head was assigned an imaging position.

Bright-field and sequential fluorescence z-stacks were then acquired at each position. The central plane of each tumor was set manually, followed by automated z-stack acquisition with a 7.5 µm step size over a total range of 130 µm using the 20× objective. Fluorescence was detected using the corresponding filter sets for LA-eGFP, mCherryNuc6, and LA-iRFP. For longitudinal experiments, the same larvae were re-anesthetized, repositioned in the molds, and imaged again under the same acquisition settings at the indicated time points. At the end of the experiments, zebrafish larvae were euthanized by an overdose (400mg/L) of MS-222, followed by decapitation with a scalpel to ensure death.

### Tumor image processing and morphometric analysis

Three-dimensional image stacks were converted into two-dimensional representations for quantitative analysis. Each optical slice was smoothed using a Gaussian filter, followed by log transformation and maximum-intensity projection. Background subtraction was performed on the resulting two-dimensional projection. Xenograft regions were identified using a ResNet50-based segmentation model trained on 200 manually segmented tumor images. All segmentations were visually inspected and manually corrected when necessary. From each segmented tumor, area, perimeter, compactness (perimeter²/area), convex hull area, and fractal dimension were extracted to quantify changes in tumor size, shape, spatial extent, and morphological complexity. Fractal dimension was calculated as described by [40].

### Immunofluorescence imaging and quantification

Whole-mount immunofluorescence was imaged using a Leica HCS LSI laser-scanning confocal microscope equipped with a 40×/1.15 oil-immersion objective and controlled using LAS AF software (Leica Microsystems). Images were acquired at 1024 × 1024 pixels using bidirectional scanning. Z-stacks were acquired at 0.76-µm intervals, with the z-range adjusted for each sample to encompass the entire xenograft. Laser power, detector gain, offset, and other acquisition parameters were kept constant within each staining experiment.

Images were analyzed in Fiji (ImageJ). Maximum-intensity projections were generated from the z-stacks, and the tumor region was manually delineated as a region of interest (ROI). Mean fluorescence intensity within the tumor ROI and background fluorescence from an adjacent non-fluorescent region were measured. Background-corrected fluorescence intensity was calculated for each xenograft and used to quantify Ki67 and active Caspase-3 staining.

Statistical analyses were performed using GraphPad Prism 10. Comparisons between two groups were performed using the Mann–Whitney U test. Experiments containing three groups were analyzed using the Kruskal–Wallis test followed, when significant, by Dunn’s multiple-comparisons test. Statistical significance was defined as p < 0.05.

### Whole-mount imaging of tumor–microenvironment interactions

To visualize interactions between xenografted tumor cells and the brain microenvironment, larvae carrying fluorescent vascular or macrophage/microglial reporter transgenes were collected at the indicated time points and fixed in 4% PFA for 2 h. Larvae were washed four times for 5 min in PBS and transferred sequentially through 25%, 50%, and 70% glycerol in PDT (PBS containing 1% BSA and 1% Triton X-100) for optical clearing.

Whole-mount larvae were imaged using a BC43 confocal microscope. Z-stacks encompassing the entire head were acquired with a 40× objective to capture the cerebral vasculature and macrophage/microglial populations in relation to the xenograft. Image stacks were acquired with a 7-µm z-step. Vascular networks were reconstructed in Imaris using manual thresholding, whereas mpeg-positive macrophages/microglia were manually quantified in Fiji. For macrophage/microglial analysis, both the total number of mpeg-positive cells within the brain and the number located within or in direct contact with the tumor were quantified.

### Statistical analysis

Statistical analysis was performed using GraphPad Prism 9.3.1 software. For morphometric images, ANOVA one-way was used to assess the effects of compounds in different body structures. For xenotransplants, mixed effect model for repeated measures was used, with Tukey’s test for multiple comparisons. Results with p-values ≤ 0.05 were considered as significant (* = P < 0.05, ** = P < 0.01, *** = P < 0.001, **** = P <0.0001, ns= not significant).

## Supporting information

Three-dimensional reconstruction of ONS-76 tumor cells and the cerebral vasculature.

## Data availability

Raw images and associated data are stored on secure servers at the University of Zurich and are available from the corresponding author upon reasonable request. The analysis scripts used in this study are publicly available in GitHub in https://github.com/veroakle/ZebraXenoSeg.

## Authors Contributions

V.A.: Conceptualization, Methodology, Investigation, Formal analysis, Data curation, Visualization, Supervision, Writing: original draft, review & editing.

A.A.: Methodology, Software, Formal analysis, Data curation, Visualization. B.K.: Methodology, Investigation.

S.S.: Methodology, Investigation.

J.D.: Methodology, Resources, Investigation. M.T.S.: Investigation, Resources.

B.C.: Software, Methodology, Resources, Investigation.

M.B.: Conceptualization, Methodology, Formal analysis, Funding acquisition, Supervision, Writing – review & editing.

S.C.F.N.: Conceptualization, Methodology, Formal analysis, Funding acquisition, Supervision, Writing – review & editing.

## Ethics Statement

All animal experiments were conducted in accordance with applicable institutional and national regulations. All protocols and experimental procedures involving zebrafish were approved by the Cantonal Veterinary Office, Zürich (Kantonales Veterinäramt), under licenses TV4206 and ZH073/2024. The Cantonal Commission for Animal Experimentation serves as the independent ethical review body, while the University of Zurich oversees institutional compliance through its Animal Welfare Officers.

## Acknowledgements

We would like to thank Matthias Gesemann for his support with animal licensing, Kara Kristiansen and Martin Walther for expert animal care and administration, Heiko Gäthje (Evident) for developing the neural-network model used for sample detection and Julian Arevalo for his support and assistance with manuscript preparation.

