## Supplemental Figures for "Quantitative live imaging reveals region-dependent medulloblastoma dissemination in the zebrafish brain"

Supplementary material

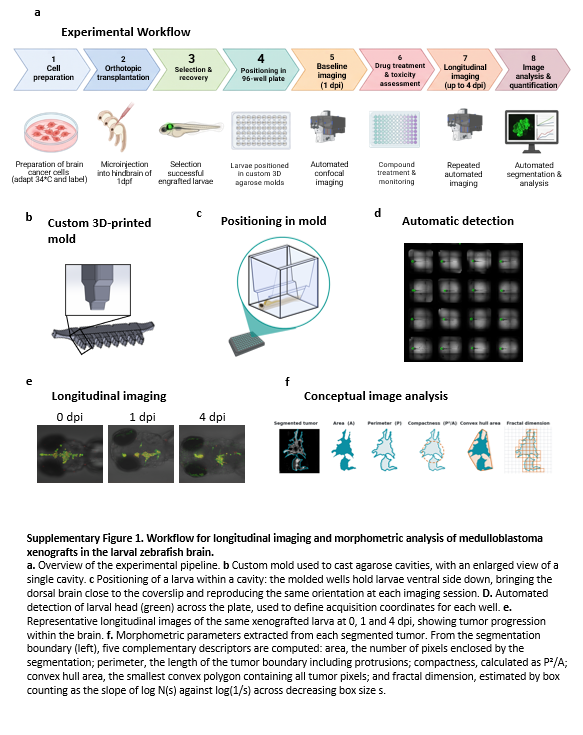

Supplementary Figure 1. Workflow for longitudinal imaging and morphometric analysis of medulloblastoma xenografts in the larval zebrafish brain.

a. Overview of the experimental pipeline. b Custom mold used to cast agarose cavities, with an enlarged view of a single cavity. c Positioning of a larva within a cavity: the molded wells hold larvae dorsal side down, bringing the dorsal brain close to the coverslip and reproducing the same orientation at each imaging session. D. Automated detection of larval head (green) across the plate, used to define acquisition coordinates for each well. e. Representative longitudinal images of the same xenografted larva at 0, 1 and 4 dpi, showing tumor progression within the brain. f. Morphometric parameters extracted from each segmented tumor. From the segmentation boundary (left), five complementary descriptors are computed: area, the number of pixels enclosed by the segmentation; perimeter, the length of the tumor boundary including protrusions; compactness, calculated as P²/A; convex hull area, the smallest convex polygon containing all tumor pixels; and fractal dimension, estimated by box counting as the slope of log N(s) against log(1/s) across decreasing box size s.

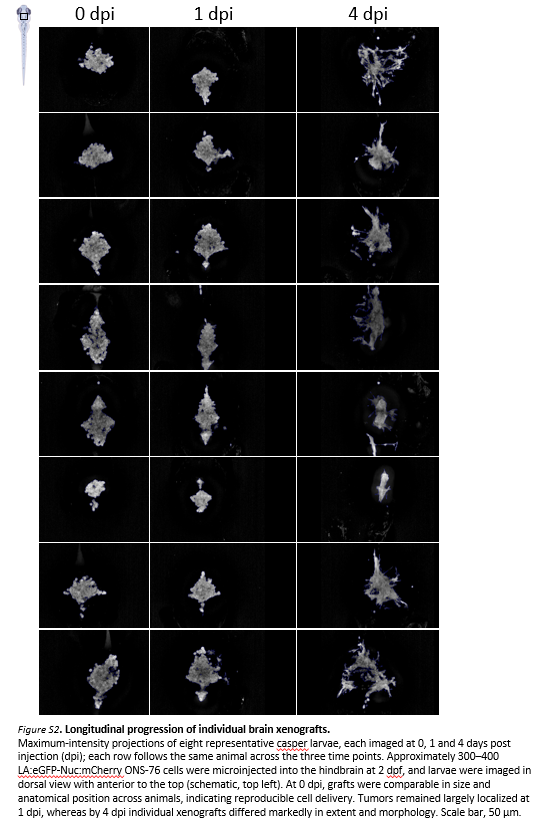

Figure S2. Longitudinal progression of individual brain xenografts.

Maximum-intensity projections of eight representative casper larvae, each imaged at 0, 1 and 4 days post injection (dpi); each row follows the same animal across the three time points. Approximately 300–400 LA:eGFP-Nuc:mCherry ONS-76 cells were microinjected into the hindbrain at 2 dpf, and larvae were imaged in dorsal view with anterior to the top (schematic, top left). At 0 dpi, grafts were comparable in size and anatomical position across animals, indicating reproducible cell delivery. Tumors remained largely localized at 1 dpi, whereas by 4 dpi individual xenografts differed markedly in extent and morphology. Scale bar, 50 µm.

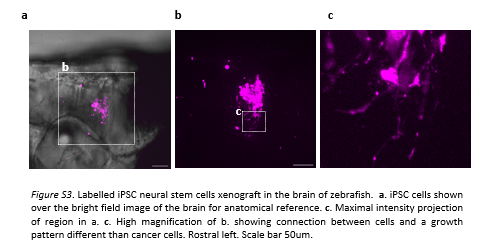

Figure S3. Labelled iPSC neural stem cells xenograft in the brain of zebrafish. a. iPSC cells shown over the bright field image of the brain for anatomical reference. c. Maximal intensity projection of region in a. c. High magnification of b. showing connection between cells and a growth pattern different than cancer cells. Rostral left. Scale bar 50um.

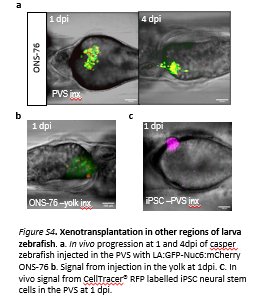

Figure S3. Xenotransplantation in other regions of larva zebrafish. a. In vivo progression at 1 and 4dpi of casper zebrafish injected in the PVS with LA:GFP-Nuc6:mCherry ONS-76 b. Signal from injection in the yolk at 1dpi. C. In vivo signal from CellTracer® RFP labelled iPSC neural stem cells in the PVS at 1 dpi.

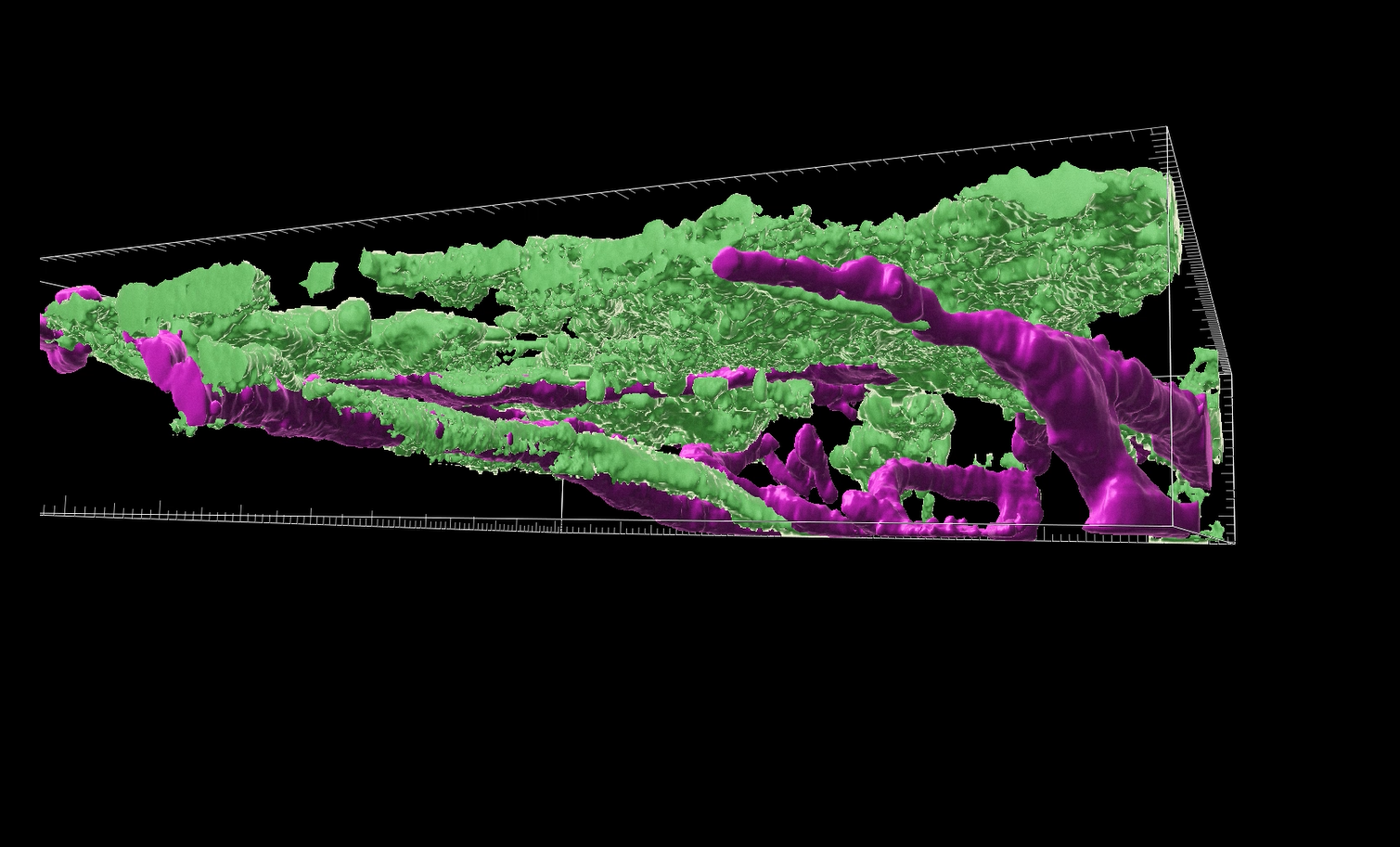

Video S1 (still)

Video S1. Three-dimensional reconstruction of ONS-76 tumor cells and the cerebral vasculature. Transgenic vascular endothelium is shown in magenta and ONS-76 tumor cells in green. Tumor cells extend along and around blood vessels during dissemination. Digital removal of the vascular signal reveals indentations in the tumor surface corresponding to the position of the vessels, further illustrating the close spatial association between tumor cells and the vasculature.

Table S1. Mixed-effects analysis of dasatinib-treated ONS-76 xenografts. Mixed-effects models (REML) were used to assess the effects of time, treatment, and their interaction on tumor morphometric parameters. P < 0.05 was considered statistically significant.

| **Morphometric parameter** | **Time (1 vs. 4 dpi)** | **P value** | **Treatment (5 groups)** | **P value** | **Time × Treatment** | **P value** |
| --- | --- | --- | --- | --- | --- | --- |
| **Area** | F(1, 80) = 2.40 | 0.1250 | F(4, 138) = 4.86 | **0.0011** | F(4, 80) = 4.35 | **0.0031** |
| **Perimeter** | F(1, 80) = 56.6 | **<0.0001** | F(4, 138) = 4.56 | **0.0017** | F(4, 80) = 3.27 | **0.0155** |
| **Compactness** | F(1, 80) = 21.3 | **<0.0001** | F(4, 138) = 0.551 | 0.6986 | F(4, 80) = 0.499 | 0.7365 |
| **Fractal dimension** | F(1, 80) = 16.6 | **0.0001** | F(4, 138) = 3.26 | **0.0138** | F(4, 80) = 1.01 | 0.4088 |
| **Convex hull area** | F(1, 80) = 42.8 | **<0.0001** | F(4, 138) = 3.71 | **0.0067** | F(4, 80) = 4.15 | **0.0041** |
| **Model details** | | | | | | |
| **Model** | Mixed-effects model (REML); fixed effects tested with Type III tests. | | | |  |  |
| **Factors** | Time = row factor (2 times); Treatment = column factor (5 levels: E3, DMSO, dasatinib 5, 10 & 15 µM). | | | | | |
| **Repeated structure** | Subject included as the matched/repeated random effect. | | | |  |  |
| **Sample summary** | 143 subjects; 58 missing values in each exported analysis. | | | |  |  |
| **Inference** | α = 0.05; sphericity assumed in the Prism analyses. | | |  |  |  |
| **Reporting** | F statistics are shown as F(numerator df, denominator df). No post-hoc comparisons are included. | | | | | |
